# Axonal ensembles repeatedly cluster and order synapses along dendrites in mouse cortex

**DOI:** 10.64898/2026.03.31.715753

**Authors:** Saarthak Sarup, Kwabena Boahen

## Abstract

Neuronal ensembles—groups of neurons that exhibit coordinated activity during behavior—are a fundamental feature of cortical computation. Dendritic branches amplify clustered synaptic inputs through local nonlinearities, suggesting that presynaptic groups might organize their connections in specific spatial patterns to engage these mechanisms. Whether the same axon groups form synaptic clusters with consistent spatial arrangements across different target neurons remains unknown, but nanoscale connectomes would resolve such anatomical motifs if they exist. We analyzed millions of synaptic connections in a connectome of mouse visual cortex and found over 700,000 axon groups that repeatedly cluster their synapses onto dendritic branches of multiple pyramidal cells, with over 500,000 maintaining consistent distal-to-proximal arrangements. These repeated patterns occur far more frequently than expected from spatial proximity or layer-based connectivity rules. Axon groups preferentially target specific dendritic branches and position their synapses in stereotyped spatial configurations across multiple postsynaptic partners, revealing that functional ensembles leave characteristic anatomical signatures in cortical microarchitecture.

## Axodendritic motifs of cortical wiring

Cortical circuits process information through coordinated activity of neuronal ensembles, with specific groups of neurons firing together during sensory experience, motor behavior, and memory formation. In the rodent hippocampus, about a dozen place cells communicate a short spatial trajectory to the cortex—a *replay*—by spiking in a particular temporal sequence during brief periods of synchrony (sharp-wave ripples) [1,2,3,21]. Multiple ensembles reactivate in sequence, chaining together short trajectories to communicate the rodent’s full path [4]. Similarly, in the rodent visual cortex, groups of ten to twenty neurons spike in synchrony, either spontaneously or in response to visual stimuli, and these ensembles activate in a repeatable sequence to communicate changing visual percepts [5,6]. For this coordinated ensemble activity to influence a downstream neuron, its dendrites must integrate inputs from multiple members of the presynaptic ensemble.

A pyramidal neuron amplifies synaptic inputs that arrive together on the same dendritic branch, with co-active inputs within a short stretch triggering a local dendritic spike that boosts their influence on the neuron’s output. In the rodent cortex, uncaging glutamate onto a cluster of spines evokes a local calcium *hotspot* in a 20 to 40-μm-long segment of a basal dendrite of a Layer 5 pyramidal cell, consistent with a supralinear response [22,23]. And activating 8 spines in distal-to-proximal order along an 80-μm-long basal dendrite elicits a larger voltage at the soma than doing so in proximal-to-distal order [7]. In Layer CA1 of the rodent hippocampus, neighboring spines spaced 3 to 5-μm apart activate sequentially along dendrites of pyramidal cells during sharp-wave ripples, either towards or away from the soma to produce differential calcium transients [8]. These findings raise the possibility that an ensemble’s coordinated activity clusters its axons’ synapses along a stretch of dendrite.

An ensemble’s coordinated spiking activity could cluster its axons’ synapses along a dendrite through activity-driven spinogenesis: a spine stabilizes when activated synchronously with its neighbors, and a dendrite extends a filopodium toward a nearby axon that fires together with existing synaptic partners. In the developing rodent cortex, a pyramidal-cell dendrite expresses NMDA but not AMPA receptors at most of its synapses [24]. These *silent* synapses contribute to depolarizing a dendrite only when neighboring afferents are co-active [34]. This in-sync activity stabilizes a silent synapse [15,24], possibly by triggering AMPA receptor insertion to unsilence it [27]. In the mature rodent cortex, a dendrite extends a filopodium—the equivalent of a silent synapse [28]—to sample a nearby axon. If its sampled axon is co-active with a nearby synaptic cluster, then the filopodium matures into a stable synapse [29]. Through this spinogenesis, coordinated spiking activity could cluster synapses along a dendrite, but whether the same group of axons forms similar clusters on multiple target dendrites remains unknown.

If a synaptic cluster reflects a specific ensemble’s coordinated activity, then that ensemble’s axons would form clusters with the same spatial arrangement on multiple dendrites, creating a repeated anatomical motif that identifies the presynaptic ensemble. These repeated synaptic arrangements could also arise from anatomical rules. Axons might cluster on nearby dendrites merely due to spatial proximity, or neurons in the same cortical layer might share stereotyped connectivity patterns. A nanoscale connectome provides the resolution needed to distinguish ensemble-specific organization from these anatomical constraints by tracing each axon to each dendrite across a population of neurons. Here we analyze a connectome of mouse visual cortex [9] to identify groups of axons that repeatedly cluster their synapses along dendrites of multiple pyramidal cells and to find any consistent spatial arrangements. The repeated motifs we uncover occur far more frequently than predicted by proximity or layer identity, revealing that ensembles of coordinated neurons imprint their functional organization in the microanatomy of cortical circuits.

### Repeated synaptic clusters are an abundant axodendritic motif

Grouping connections between 48,000 axons and 1.5 million dendritic branches in a cubic millimeter of mouse visual cortex [9] (Fig. 1A) revealed over 0.7 million groups of axons that cluster their synapses along individual branches of multiple dendritic trees. Of these groups, over 0.5 million replicate the same distal-to-proximal arrangement (ordered cluster) (Fig. 1B). These synaptic clusters occur throughout the cortical volume, are concentrated in Layers 4 and 5, and slightly prefer basal over apical dendrites (Fig. 1C and Supp. Fig. 1).

**Figure 1:**
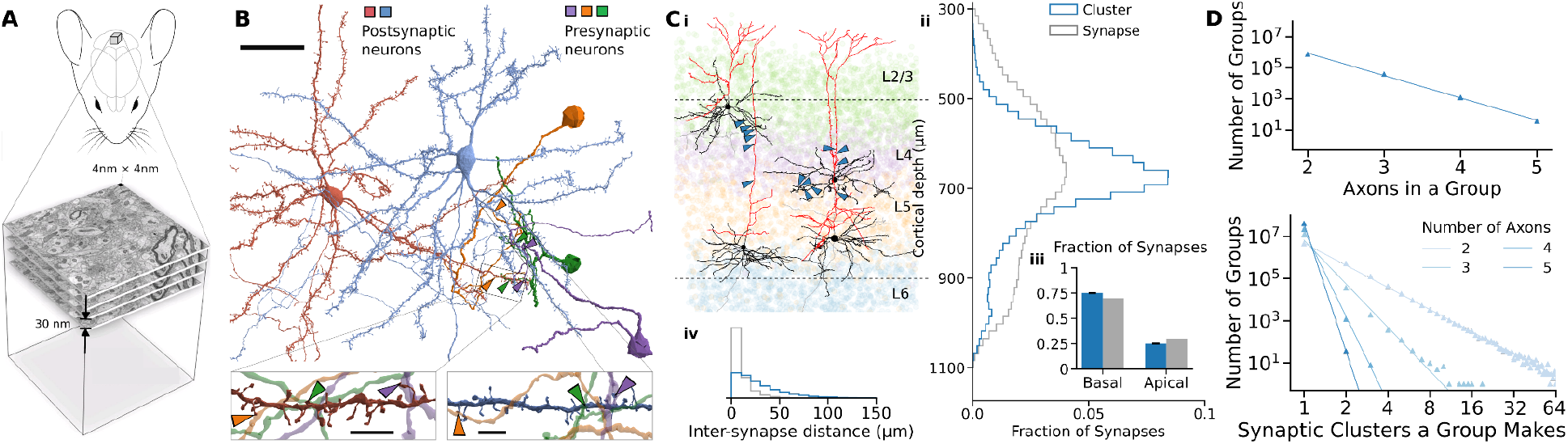
Groups of axons cluster their synapses along a dendritic branch. A – Mouse visual cortex: The MICrONS connectome identifies 48,000 pyramidal cells interconnected by 4 million excitatory synapses of 75,000 neurons and 500 million synapses reconstructed from electron-microscope images (0.93 mm^3^ *in vivo*) [18]. B – A repeated synaptic cluster: Three axons (*purple, orange, green*) synapse onto dendrites of two pyramidal cells (*red, blue*) in the same distal-to-proximal arrangement (*purple*-*green-orange*, insets). Meshes of neurons are pruned for visual clarity. Scalebar: 20 (main panel) and 5 (insets) μm. C – Distribution across cortical layers: *i*, Left: Three axons synapse (*blue*) onto a Layer-2/3 cell’s basal dendrite (*black*) and a Layer-5 cell’s apical dendrite (*red*) in different arrangements. Right: Four axons synapse (*blue*) onto a Layer-4 cell’s basal dendrite (*black*) and a Layer-5 cell’s apical dendrite (*red*) in the same arrangement. *ii*, Synapses within clusters peak (*blue*) at a similar cortical depth as all pyramidal-pyramidal synapses (*gray*). *iii*, They occur slightly more frequently on basal dendrites than expected. Whisker: 95% credible interval. *iv*, Axons forming clusters space their synapses further apart than all identified excitatory axons (median: 22 vs. 6.0 μm). D – Distribution of axon group sizes and synaptic cluster counts: *Top*, The number of groups with two or more clusters decreases with a fixed geometric ratio. *Bottom*, The number of clusters per group decreases as this group’s size increases.

Growing axons may first form ensembles and then branch together to cluster their synapses on nearby dendritic branches. If a *k*-th axon ensembles with *k* – 1 axons (*k* > 2) with probability *p*, then the number of *k*-axon groups that form repeating synaptic clusters follows a geometric distribution, with *T*(*k*) = *T*(2)*p*^*k*-2^. For groups of 2 to 5 axons, this probability was estimated as *p* = 10^-1.44^ = 1/28 (Fig. 1D, *top*), though this is likely underestimated due to sparse synapse identification (only 1 in 125). Once formed, these ensembles may branch stochastically to form synaptic clusters, resulting in a power-law distribution *N*(*c*) = *N*(1)*c*^-α^ for the number of axon groups forming *c* synaptic clusters [10,11]. As group size increases from 2 to 5 axons, the exponent *α* increases (steeper slope on log-log plot) from 3.5 to 20.1 (Fig. 1D, *bottom*), reflecting the decreasing likelihood that all axons in a group branch together at a given point. While these distributions suggest coordinated targeting, it is also possible that axons cluster their synapses simply because their arbors pass near the same dendritic branches in the dense cortical neuropil.

A presynaptic axon is often close to multiple branches of each postsynaptic dendritic tree, which creates opportunities for synaptic clusters to form even without active coordination. For example, a 5-μm distance encompasses bouton-plus-spine length for 97.5% of synapses, and within this distance, a presynaptic axon is proximal to three branches of a postsynaptic dendritic tree (median, Supp. Fig. 2A), including the branch it actually synapses on. If an axon rearranges its synapses among these proximal dendritic branches, it could form new synaptic clusters with other presynaptic axons without increasing its overall cable length (Supp. Fig. 2B). This raises the question: could the observed abundance of synaptic clusters arise simply from the geometry of axonal and dendritic arbors, if axons choose among the proximal branches at random?

**Figure 2:**
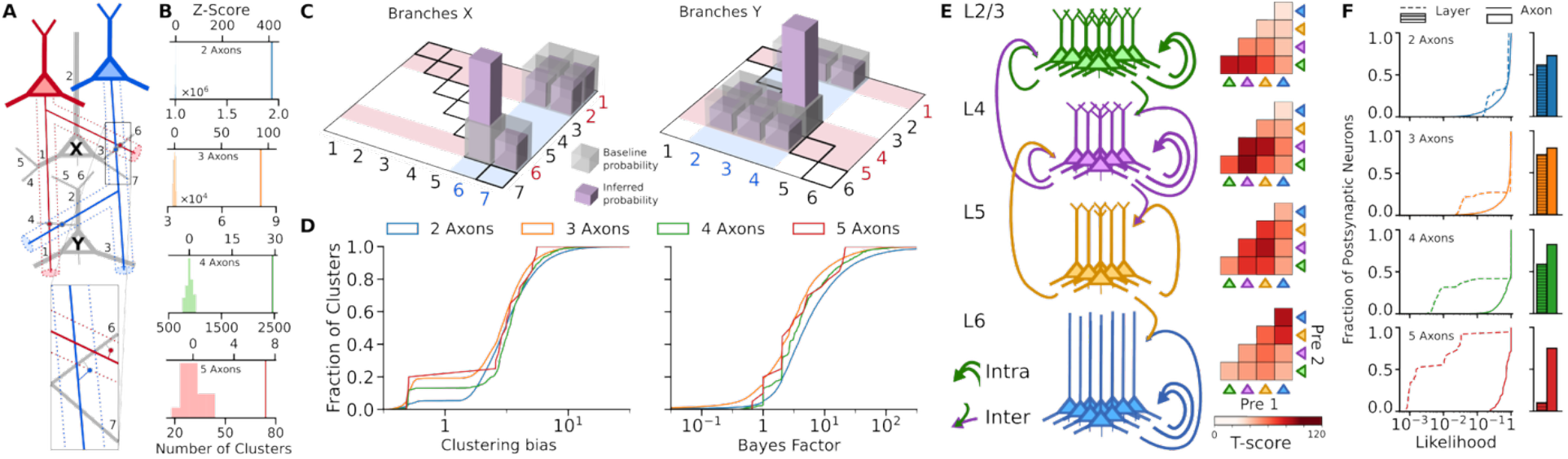
Axons that cluster span more cortical layers than pairwise attraction predicts. A – Synaptic clustering: Axons of two neurons (*red* and *blue*) are proximal (< 5 μm) to dendrites of two other neurons (X and Y) and cluster their synapses on the same branch of both (branch 6 of X, *inset*, and branch 4 of Y). B – Repeated synaptic clusters in the connectome vs. proximal shuffles (*n* = 50): The number of clusters varies from 1,934,374 vs. 1,002,790 ± 2,259 (Z = 412) for 2-axon groups to 74 vs. 31.0 ± 6.1 (7.1) for 5-axon groups. C – Clustering bias: Pairings of X’s (*left*) or Y’s (*right*) branches proximal to one or both axons (*red* and *blue* stripes intersect) set a uniform probability baseline (*gray bars*). Probabilities inferred from observing two synaptic clusters (*purple bars*) yield a clustering bias of 5.3 (ratio of *purple* to *gray bars* along diagonal). D – Cumulative distribution of clustering bias and Bayes factor: For groups of 2 to 5 axons, 95 to 76% respectively were attracted (bias > 1) to synapse on the same branch (*left*), and 65 to 43% substantially prefer (Bayes factor > 10^1/2^) doing so (*right*). E – Axon-pairing in the cortical column: *Left*, Dendrites in Layer 2/3 (*green*), 4 (*purple*), and 6 (*blue*) are mostly postsynaptic to pairs of axons from their own layer (*intra*) and from neighboring layers (*inter*) but dendrites in Layer 5 (*yellow*) prefer the latter. *Right*, T-score between inferred and baseline clustering probability (degrees-of-freedom > 143). F – Cumulative distribution of an axodendritic motif’s likelihood (*left*) and accurately predicting its occurrence (*right*) based on layer- and axon-wise preferences. Median likelihood ranges from 0.80 and 0.86 for 2-axon groups to 0.0019 and 0.74 for 5-axon groups, respectively. Prediction accuracy ranges from 61% and 73% to 10% and 75% of motifs, respectively.

After controlling for axon-dendrite proximity (Fig. 2A), we found that repeated synaptic clusters formed by groups of 2 to 5 axons are significantly more common in the cortex than would be expected by chance. Specifically, the connectome contains 1,470,000, 120,000, 4,900, and 185 repeated synaptic clusters respectively from groups of 2, 3, 4, and 5 axons—corresponding to 433, 99, 29, and 7.1 standard deviations above the mean of 50 proximal shuffle controls (Fig. 2B). This over-representation persists even when controlling for imperfections in automatically reconstructed axons (Supp. Fig. 3), such as erroneously merged neurites that misattribute a synapse’s presynaptic cell [12]. These findings indicate that axonal ensembles actively coordinate their dendritic targeting, rather than clustering through proximity alone, prompting further investigation into the mechanisms that guide this coordination.

**Figure 3:**
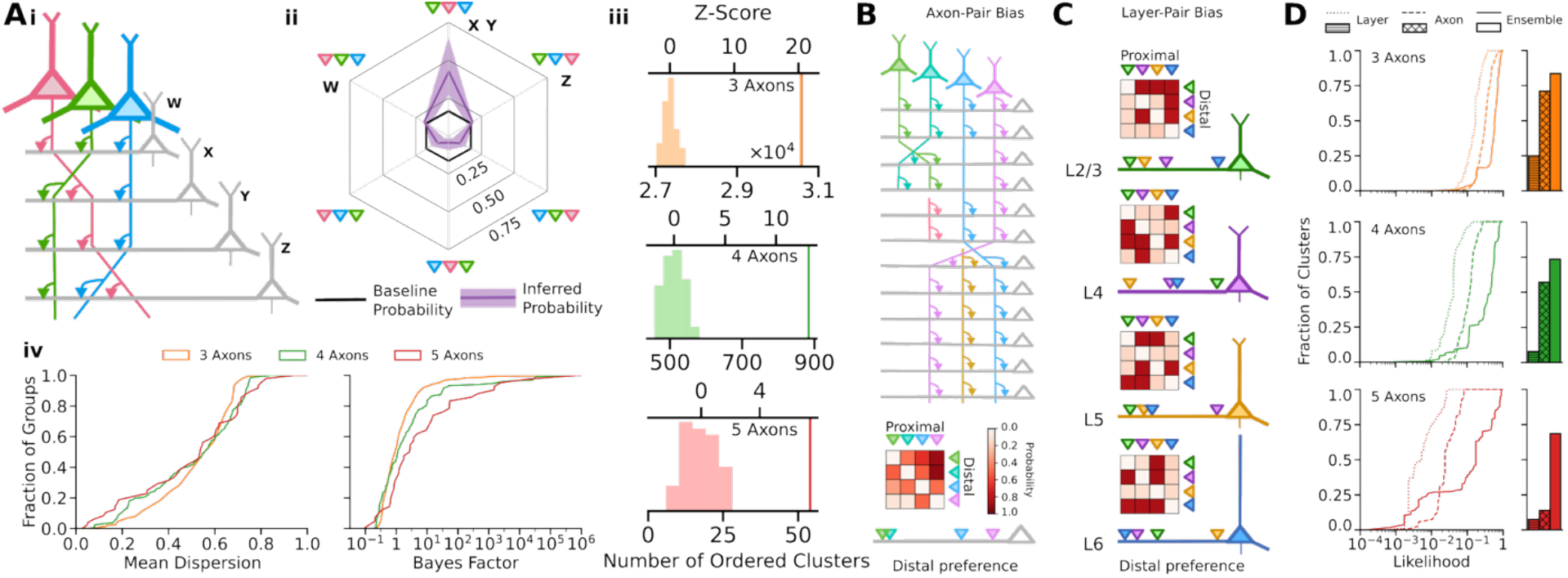
Axons often do not choose the synaptic order that their layer prescribes. A – Dispersion of synaptic orderings: *i*, Axons of three neurons (*pink, green, blue*) cluster their synapses on dendrites of four neurons (W, X, Y and Z) in three different orderings. *ii*, A Mallows distribution fit to these observations (*purple*) infers a central permutation (*green-pink-blue*) and a dispersion (mean ± S.D. = 0.452 ± 0.252). *Black*: uniform baseline. *iii*, Repeated synaptic clusters with the same distal-to-proximal order in the connectome vs. in-branch shuffles (*n* = 50) range from 30,580 vs. 27,349 ± 158 (Z = 20) for 3-axon groups to 54 vs. 17.7 ± 4.87 (7.5) for 5-axon groups. *iv*, Cumulative distribution of mean dispersion (*left*) and the Bayes factor (*right*). B – Order preference of axon-pairs: Four axons (*green-teal-blue-purple*) cluster 2 to 4 synapses along 12 dendrites, 8 of which receive a third synapse from two other axons (*pink* and *yellow*). All these axodendritic motifs yield axon-pair orderings (*matrix*) that rank the 4 axons (*bottom branch*) and predict their synaptic clusters’ central permutation correctly. C – Order preference of layer-pairs: Order preference (*matrix*) and distal preference (*branch*) for dendrites in Layers 2/3, 4, 5, and 6 (*first* to *fourth row*). See B for probability legend. D – Cumulative distribution of an axodendritic permutation’s likelihood and accurately predicting its ordering based on layer-, axon-, and ensemble-wise preferences: Medians of likelihood distributions (*left*) range from 0.17, 0.32, and 0.58 for 3-axon groups to 0.0052, 0.024, and 0.17 for 5-axon groups, respectively, and prediction accuracies (*right*) range from 25, 71, and 84% to 7.5, 14, and 69%.

### An ensemble’s axons are biased to synapse on the same branch

To assess an axon’s preference to cluster its synapse with another axon’s synapse on a branch of a dendrite, we inferred each axon-pair’s clustering bias. With a median of 3 proximal branches per presynaptic axon, there are 9 pairings of these 3 branches that 2 presynaptic axons could choose. We compare the likelihood of this axon-pair’s choices across multiple dendritic trees under two Bayesian models. Both have a latent random variable—the pair’s clustering bias—that sets the probability of synapsing on the same branch relative to the uniform baseline probability (Methods). A priori, this clustering bias is distributed between zero and unity for one model (repulsion) and between 1 and infinity for the other model (attraction). The ratio of the observations’ likelihoods (Bayes factor) under these two models yields substantial (≥ 10^1/2^) or strong (≥ 10) evidence of attraction or repulsion. For a group of *k* > 2 axons, we evaluate *k* Bayes factors for attraction and repulsion between a held-out axon and the remaining *k* − 1 axons, each with its own posterior clustering bias, and assign the most extreme (largest or smallest) Bayes factor and its corresponding bias to the group (Methods).

For example, consider two axons proximal to two dendritic trees (see Fig. 2A). The first axon is proximal to 2 branches of the first tree and 3 branches of the second tree; the second axon is proximal to 3 branches of both trees. Thus the probability of clustering on the same branch is 1/(2×3) = 1/6 for the first tree, 1/3×3=1/9 for the second tree, and 1/(6×9) = 0.0185 for both trees (Fig. 2C). Observing these two clusters yields a posterior clustering-bias distribution with mean 5.3 under the attracting model that skews the probability of clustering on the first tree from 1/6 to 1/1.9, on the second tree from 1/9 to 1/2.5, and on both from 0.0185 to 1/(1.9×2.5) = 0.21—the full distribution gives 0.24. In contrast, under the repelling model, these observations yield a mean posterior clustering-bias of 0.66 that skews the clustering probabilities from 1/6 to 1/8.6, 1/9 to 1/13.1, and 0.0185 to 1/(8.6×13.1) = 0.0089—the full distribution gives 0.0098. The ratio of these likelihoods (0.24/0.0098 = 24.5) yields strong evidence (Bayes factor > 10) for attraction.

Out of the 810,901 groups of 2 to 5 axons observed to cluster on dendrites of two or more postsynaptic neurons, our Bayesian models identified 518,677 groups that are substantially attracted (Bayes factor ≥ 10^1/2^) to synapse on a branch proximal to all of their axons and only 10,870 groups that are substantially repelled (Bayes factor ≤ 10^-1/2^) (Fig 2D). On average, a group of axons in the attracting class clustered onto the same dendritic branch in 89% of the neurons it targeted, suggesting that these grouped axons truly belong to the same ensemble. But this fraction was only 13% for the repelling class, suggesting that these grouped axons belong to distinct ensembles that just happened to synapse on the same branch. Could this axonal attraction and repulsion be explained by the cortical layer of origin?

### Axons ensemble across the cortical column

To assess the cortical layer’s role in synaptic clustering, we aggregated axon-wise probabilities across axon-pairs from and a dendrite in a set of cortical layers to obtain a layer-wise clustering probability. For instance, given axons from Layers 4 and 5 and dendrites in Layer 2/3, we average the axon-wise clustering probabilities for every Layer 4-5 axon-pair proximal to a Layer 2/3 dendrite. For groups of more than two axons (*k* > 2), we multiply *k* − 1 layer-wise clustering probabilities between the held-out axon with the most extreme Bayes factor (as described above) and each of these *k* − 1 axons. For instance, if a 3-axon group from Layers 2/3, 4, and 5 forms a synaptic cluster on a branch of a dendrite in Layer 2/3 and the Layer 5 axon is held-out, then multiplying layer-wise clustering probabilities between a Layer 5-2/3 axon-pair onto a Layer 2/3 dendrite and a Layer 5-4 axon-pair onto a Layer 2/3 dendrite yields the layer-wise clustering probability of this 3-axon group on the Layer 2/3 dendrite.

Cortical layer predicts whether a pair of axons is attracted to or repelled from synapsing on a branch of a dendritic tree better than chance, but this accuracy worsens for larger axon groups. Layer-wise clustering probabilities show that groups are most likely to include pairs of axons from the same or neighboring cortical layers (Fig. 2E). For 2-axon groups, these probabilities correctly predict clustering on 61% of dendritic trees postsynaptic to both axons—above the 50% correct expected by chance (Fig. 2F, *1st row*). Prediction accuracy rises to 72% for 3-axon groups but then drops to 59% for 4-axon groups and to 10% for 5-axon groups (Fig. 2F, *2nd to 4th row*). Notably, among these incorrect predictions, 25%, 41%, and 67% of these respective groups span three or more cortical layers. In contrast, predictions based on each group’s clustering bias remain consistently accurate—between 73% and 82% correct—regardless of group size. Thus, the consistent clustering of multilayer axon groups reflects ensemble-specific coordination rather than layer-specific rules, leading to the question of whether this coordination also governs the spatial ordering of synapses within each cluster.

### Axodendritic combinations and permutations

To assess whether axon groups maintain consistent spatial ordering of their synapses or position them randomly along dendritic branches, we model each group’s observed synaptic orderings using a Mallows distribution (Fig. 3A, *i & ii*) [13]. This probability distribution is parameterized by a *central permutation* and a *dispersion*—the probability of swapping adjacent elements—0 corresponds to maximal order and 1 corresponds to maximal disorder. That is, only the preferred permutation occurs and all orderings occur with equal probability, respectively. For example, a group of 3, 4, or 5 axons may form orderings that differ by up to 3, 6, or 10 swaps. When fit with a dispersion of 0.5, their synaptic orderings differ by an average of 1.25, 2.33, or 3.6 swaps, respectively.

Controlling for dendritic branch targeting, synaptic clusters formed by groups of 3 to 5 axons that repeat in the same distal-to-proximal order on two or more dendritic trees are over-represented in the cortex. We shuffled the order of axons’ synapses along their targeted dendritic branch while preserving the connectome’s synaptic clusters. The connectome has 30,000, 880, and 54 ordered clusters respectively from groups of 3, 4, and 5 axons—20 to 5.4 standard deviations more than the average of 50 within-branch shuffles (Fig. 3A, *iii*). Axon groups therefore do not position their synapses randomly along dendritic branches but rather synapse with small dispersion about a preferred ordering.

To identify axon groups with substantial evidence of order preference, we compared the likelihood of each group’s observed synaptic orderings under two competing Bayesian models. Both model these orderings as samples from a Mallows distribution but differ in their prior distribution of dispersion. In one model, the dispersion concentrates at 1, reflecting a fixed belief that all permutations of a group’s axons are equally likely. In the other, the dispersion is sampled from a uniform prior distribution ranging from 0 to 1, reflecting a subjective belief that no value of dispersion is favored. The observed permutations yield a posterior distribution of dispersions and likelihoods under each model (Fig. 3A, *iv* and Methods).

The ratio of likelihoods under these models (Bayes factor) isolated subsets of axon groups that consistently formed disordered and ordered synaptic clusters, named *axodendritic combinations* and *permutations*, respectively. For groups of 3 to 5 axons, Bayes factor comparisons isolated 2,261 axodendritic combinations (Bayes factor ≤ 10^-½^), whose synaptic orderings differ by up to 89% of the maximum number of swaps; 29,946 indecisive groups (10^-½^ < Bayes factor < 10^1/2^), whose synaptic orderings differ by up to 57% of the maximum number of swaps; and 8,101 axodendritic permutations (Bayes factor ≥ 10^1/2^), whose synaptic orderings differ by at most 18% of the maximum number of swaps. What determines their preferred synaptic ordering along a stretch of dendrite? Is it the distal-to-proximal preference of their cortical layers or of the axons themselves?

### Synaptic order within a cluster is not layer-specific but rather ensemble-specific

We used the 8,101 axodendritic permutations’ inferred central permutation and dispersion to obtain an axon’s or layer’s distal preference (axon-pair and layer-pair ordering probabilities) and thus predict the synaptic ordering within a cluster (Fig. 3B,C and Methods). Given an axon pair, we calculate the expected probability that one axon synapses closer to the soma than the other by marginalizing over the remaining axons’ positions across all axodendritic permutations with that pair. Given a layer pair, we calculate the expected probability of its distal-to-proximal ordering across all pairs of axons from that pair of layers onto dendrites in a given layer. Given either of these pairwise-order probabilities, we use the Bradley-Terry model to infer a latent *distal preference* [14] and the Plackett-Luce model to determine a distribution over all distal-to-proximal permutations [15,16] (Methods). If indeed pairwise orderings yield a distal preference that correctly predicts the observed synaptic orderings of each axodendritic permutation, then synaptic orderings would not be specific to their ensemble but rather to their cortical layers or to the pairs of axons within the cluster.

Layer-pair orderings predict a cluster’s synaptic ordering with only 25% accuracy for groups of 3 axons, and this accuracy falls to 7.6% for groups of 5 axons, largely because pairs of axons often violate their layers’ order preferences (Fig. 3D). In clusters where the order is correctly predicted, only 4.6% of axon-pairs violate their layer-pair’s order preference, whereas in clusters with incorrect predictions, 42% of axon-pairs do so. These results indicate that while cortical layer can influence the order in which some axons synapse along a dendrite, it does not reliably determine the synaptic ordering within a cluster.

Axon-pair orderings predict a cluster’s synaptic ordering with 71% accuracy for groups of 3 axons but this accuracy falls to 14% for groups of 5 axons, because the additional axons reverse a pair’s preference towards the opposite ordering (see Fig. 3D). Axon-pair orderings are most often correct when their opposite ordering is never observed in another group’s central permutation, accounting for 78% of all correct predictions. In contrast, every cluster whose ordering is incorrectly predicted contained an observation in the opposite order. In general, the average axon-pair reverses its order in 25% of incorrectly predicted orderings versus only 3% of correctly predicted ones. Thus while pairwise preferences often guide ordering among smaller groups, these preferences become unpredictive as group size increases.

While axon-pair orderings become unpredictive as groups grow in size, ensemble-specific preferences—captured by the central permutation inferred for an axodendritic permutation— remain predictive. Ensemble-specific preferences predicted the observed ordering correctly for 84%, 74%, and 69% of synaptic clusters of sizes 3, 4, and 5, respectively, far exceeding the accuracy of layer-pair and axon-pair preferences (see Fig. 3D). These results provide strong evidence for ordered dendritic input—even when axonal or layer biases suggest otherwise.

In summary, we have uncovered evidence that ensemble-specific spike sequences guide synaptic placement along dendrites in the cortex. A group of axons does not synapse onto nearby cortical dendrites randomly or according to a static blueprint imposed by cortical layer, but rather actively architects its synapses along stretches of multiple dendritic trees. These repeated axodendritic permutations are ideally situated to mediate ensemble-to-ensemble communication with a sequence-based neural code.

## Discussion

Synaptic clustering occurs far above chance levels, manifesting as 518,677 groups attracted to cluster on the same branch of 89% of their postsynaptic targets, contrasting sharply with 10,870 groups repelled to cluster on different branches of 87% of their targets. A group’s attraction or repulsion is reflected in the probability relative to random chance that it forms a cluster. This clustering bias’ predictions are consistently accurate—between 73% and 82% correct— regardless of group size. Whereas layer-based predictions decline from 61% for pairs to 10% for groups of five axons. These findings suggest that an axon group’s dendritic branch targeting does not arise from layer-wide spike patterns but rather from an ensemble-specific sequence of spikes emitted in synchronous bursts. Perhaps through an “out-of-sync, lose-your-link” developmental rule [17] (Fig. 4A).

**Figure 4:**
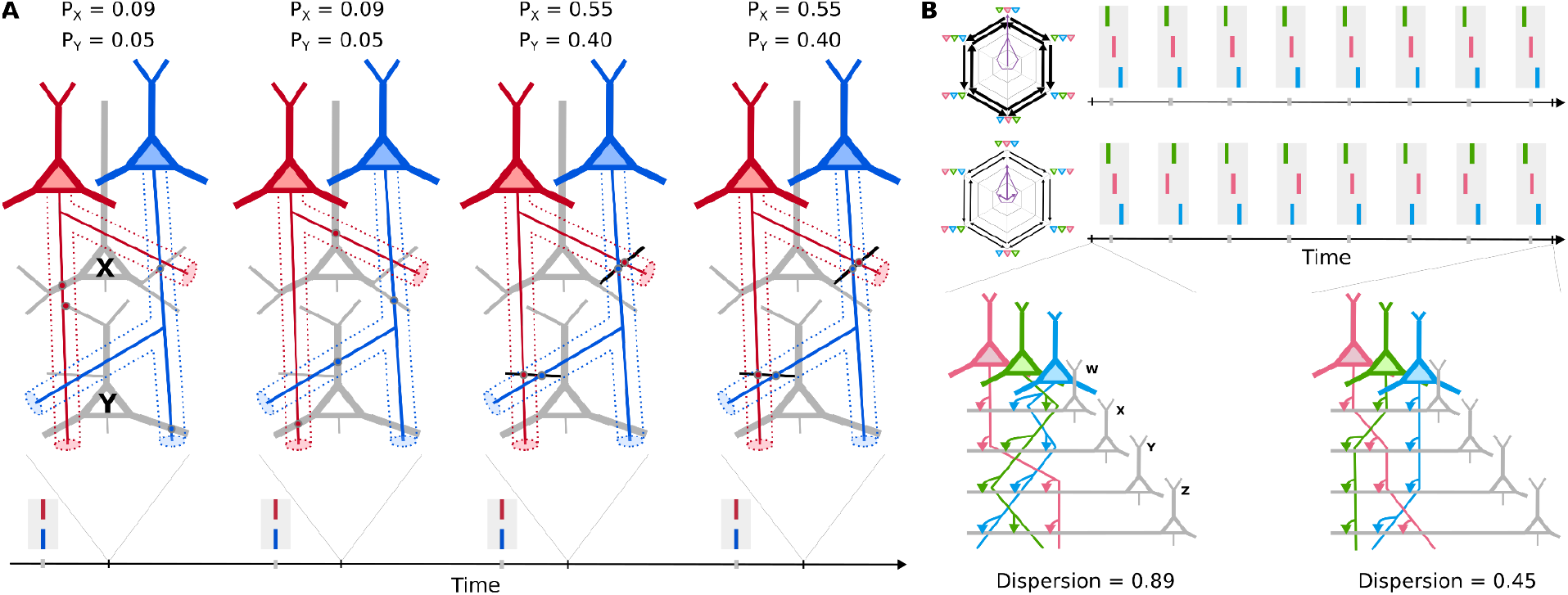
How synaptic clustering and ordering could arise. A – Synaptic clustering: Axons of two neurons (*red, blue*) synapse on dendritic branches of two neurons (X and Y). Probabilities P_X_ and P_Y_ (clustering bias = 5.3) reflect their spikes (*rasters*) temporal coherence with those branches’ other synaptic inputs. Through successive plasticity events (*left and middle-left panels*), these axons converge on the same dendritic branch (*middle-right panel*), where synaptic potentiation stabilizes synchronized inputs (*right panel)*. B – Synaptic ordering: *Top row*, An ensemble spiking in a consistent sequence (*green-pink-blue*) induces drift (*one long purple arrow*) towards that central permutation, reducing dispersion of initially disordered synaptic arrangements found across dendritic branches (*left circuit*). Frequent stochastic rearrangements (*thick black arrows*) induce diffusion towards a maximally disordered state. Balancing these forces yields the partially ordered state that we observe (*right circuit*). *Bottom row*, This equilibrium could also arise from less frequent synaptic rearrangements (*thin black arrows*) and more frequent spike reordering (*purple arrows with nonuniform length*).

Beyond clustering on the same branches, a subset of axonal ensembles also order their synapses consistently along branches of multiple dendritic trees. The dispersion of these axodendritic permutations ranges from 0.17 to 0.23, in contrast to 0.89 for the highly disordered arrangements seen in axodendritic combinations. This consistency in axodendritic permutations reflects coordination within an ensemble, as even small groups like axon pairs can prefer different orderings when they participate in different ensembles. Moreover, the central permutation predicts an axodendritic permutation’s synaptic arrangements with 69 to 84% accuracy, compared to 7 to 25% for rules based on an axon’s or dendrite’s cortical layer. These findings indicate that both clustering and ordering are driven by ensemble-specific coordination rather than layer-based developmental programs, possibly through a balance of drift (from consistent spiking order) and diffusion (from stochastic synaptic rearrangement) (Fig. 4B).

The superiority of ensemble-specific predictions over layer-based predictions—75-82% vs. 61-10% for clustering and 69-84% vs. 7-25% for ordering—suggests that coordination mechanisms operate independently of cortical layer structure. Since only 1% of excitatory synapses onto pyramidal cells were sampled, a high clustering bias could indicate either tight coordination in a small ensemble or loose coordination in a larger ensemble that sampled a coordinated subset. Similarly, low dispersion could indicate either tight ordering in a small ensemble or loose ordering in a large ensemble with shared subsequences. However, even with unbiased subsampling, the relative differences remain significant: groups with higher clustering bias coordinate more strongly than those with lower bias, and groups with lower dispersion exhibit tighter ordering than those with higher dispersion. Thus, ensemble-specific predictions’ are consistently more accurate than those based on cortical layer.

In the dense neuropil of the mouse visual cortex, where highly branched axons and dendrites offer countless potential synaptic sites, we find groups of pyramidal-cell axons that repeatedly cluster their synapses onto the same dendritic branches of multiple pyramidal cells, with most maintaining consistent spatial arrangements. The dendritic branches an axon group targets and its synaptic orderings along those branches cannot be accurately predicted by cortical layer or anatomical proximity. Instead, an ensemble’s coordinated activity could establish these axodendritic motifs through activity-dependent mechanisms, thereby sculpting these repeated patterns in the cortex’s microarchitecture independently of layer-based developmental programs. These activity-dependent mechanisms may represent a fundamental principle by which cortical circuits encode experience in the synaptic organization of dendrites.

## Acknowledgements

This study was supported by US National Science Foundation (grant number 2223827), Stanford Human-Centered Artificial Intelligence and Wu Tsai Neurosciences Institutes (Partnership Seed Grant), and Stanford Medical Center Development (Discovery Innovation Fund). We thank F. Collman for technical assistance with the MICrONS Cubic Millimeter dataset and A. Tolias for his advice and support. We also thank members of the Tolias Lab, Z. Ding and S. Papadopulous, and of the Boahen Lab, G. Weintraut and N. Reidman, for their helpful technical discussions throughout the study.

## Methods

### Data availability

All data for this study were from the MICrONS Cubic Millimeter dataset. Pyramidal cell classifications (L2/3, L4, L5-ET, L5-IT, L5-NP, L6-CT, L6-IT), cell segmentations, and synapse coordinates and pre/post segmentation IDs were accessed from version 343 of the dataset’s public release through CAVE [18] and with the API of the navis Python library [19].

### Dataset curation

We analyzed synaptic connectivity among all 48,669 neurons classified as pyramidal cells with at least one incoming synapse (dataset version 343), regardless of proofreading status, to achieve sufficient sample size for identifying repeated axodendritic motifs across populations of neurons. A neuron’s dendrite or axon is proofread when it is removed of all erroneous merges (partial/clean proofreading) and its tip is extended throughout the dataset’s volume (complete/extended proofreading). In version 343, only 751 neurons had at least partially proofread dendrites and 410 had partially proofread axons, yielding circuits too sparse for systematic motif detection. In the current dataset version (1412) that includes more extensive proofreading, 2,129 neurons have partially proofread dendrites and 2,017 have partially proofread axons. Despite this increased proofreading, connectivity among only these neurons remained sparse—2,784 synapses connected these axons and dendrites, and only 119 neurons had a dendritic branch with 2 or more synapses from these axons. Searching for axodendritic motifs among this larger proofread subset, we identified 12 pairs of neurons that repeatedly formed synaptic clusters: 11 pairs each clustered on two different dendritic branches and one pair clustered on four different branches, yielding a total of 26 synaptic clusters. These findings demonstrate that similar motifs emerge in the proofread subset despite its limited connectivity, motivating expanding this search to the full dataset.

### Finding repeated synaptic clusters

Pyramidal cells were skeletonized with the API of the navis Python library using the TEASAR algorithm into a tree rooted and directed towards the soma. Terminal branches of a skeleton with a length equal to or below 3 microns were classified as spines and pruned from the skeleton. A dendrite’s incoming excitatory synapses were assigned to their closest node in the skeleton using a KD-Tree. Each skeleton was broken at nodes with more than one predecessor into a set of branches—a chain of nodes with at most one predecessor and at most one successor, ordered distal-to-proximal towards the soma.

Every branch of one pyramidal cell was compared to every branch of every other pyramidal cell based on the set of presynaptic axons to find synaptic clusters and their distal-to-proximal order to find ordered synaptic clusters. Given two branches to compare and their ordered list of presynaptic axons, we computed the intersection of their sets. If it included two or more axons, we recorded the intersection and the pair of branches. Each recorded intersection identifies a synaptic cluster that repeats on the dendritic branches of at least two pyramidal cells. To find these clusters efficiently, we avoid exhaustively comparing all pairs of branches, which scales quadratically with the total number of branches, for a parallelized approach that scales log-linearly.

To achieve a log-linear scaling, we initialize the matrix of intersections, reorder each branch’s axons by their presynaptic cell ID, and organize the branches into a min-heap by their first (and smallest-valued) cell ID. Within this initialization, reordering each branch takes time linear in the number of branches and organizing them into a min-heap takes log-linear time. We then iteratively remove branches until the heap is empty. To do so, we remove the branch at the top of the heap (in constant time) and pop its first cell ID and record it. We continue removing the next-smallest branches and popping their first cell ID until that ID no longer matches the recorded ID. For those that were removed because their popped ID matches the recorded cell ID, we add that ID into the matrix of intersections for every pair of branches that was removed. Any removed branch that still has remaining cell IDs is re-inserted into the min-heap (in logarithmic time). The number of times this iterative step is performed scales linearly with the number of branches, and so its overall time scales log-linearly. Including the log-linear initialization time, this parallelized approach scales log-linearly with the number of branches. At its completion, the matrix of intersections has recorded the intersection between every pair of branches and has thus recorded the synaptic clusters common to at least two dendritic branches. We ignore clusters with fewer than two axons, and for those with two or more, we proceed to evaluate whether they follow a common ordering.

To find ordered synaptic clusters that repeat, we calculate all common subsequences between the distal-to-proximal orderings of every pair of branches whose synaptic clusters were formed by the same set of presynaptic axons. When a common subsequence included two or more axons, and every axon in the subsequence was unique, we recorded the subsequence and the IDs of the pair of branches. Each recorded common subsequence identifies an ordered cluster that repeats on at least two dendritic branches.

### Credible intervals

To estimate the 95% credible intervals for the fraction of synapses in unordered and ordered clusters along apical or basal dendritic compartments, we fit the counts to a Beta-Binomial Bayesian model. The Beta distribution—the model’s prior—was parameterized by the fraction of synapses found along the apical or basal compartment. Based on the number of synapses in an unordered cluster and ordered cluster that we found on the two compartments, we calculated the parameters of the posterior distribution—also Beta-distributed—and used its inverse CDF to find the mean-centered range that contains 95% of the posterior’s probability mass.

To estimate the credible intervals for the fraction of axons of a given cell-type that repeatedly formed synaptic clusters, we fit the counts to a Dirichlet-Multinomial Bayesian model. We identified the set of cells whose axons repeatedly clustered and grouped them by their defined cell-type. The Dirichlet distribution—the model’s prior—was parameterized by the fraction of cells of a given cell-type across the entire volume. Based on the number of cells of a given cell-type that form repeated clusters, we calculated the parameters of the posterior distribution (Dirchlet-distributed), marginalized out the other groups to define a Beta-distributed posterior, and used its inverse CDF to find the mean-centered range that contains 95% of this Beta distribution’s probability mass. We followed an identical procedure to estimate the credible intervals for the fraction of dendrites of a given cell-type that received synaptic clusters.

### Shuffling synapses

To randomly shuffle synapses across a pyramidal cell’s dendritic tree, we assigned each edge between nodes in the skeleton a probability proportional to the edge’s length (measured as a Euclidean distance from distal node to proximal node) and independently sampled an edge for each synapse. A synapse’s exact position along that edge was found by sampling a value from Uniform(0,1) and repositioning the synapse this fraction of the distance along the edge. After repositioning all synapses on a pyramidal cell’s skeleton, we followed our original procedure to partition the skeleton into a set of branches, but whose synapses were now redistributed and reordered along all of its branches.

To proximally shuffle synapses across a pyramidal cell’s dendritic tree, we first isolated a skeleton for each pyramidal cell’s axon and mapped its dendritic proximities. We isolated an axonal skeleton by assigning a pyramidal cell’s outgoing synapses to nodes of its complete skeleton and preserving only the nodes whose path towards the soma or whose path towards a leaf included an outgoing synapse. To efficiently check whether a point on a dendrite was within a 5-μm proximity of a presynaptic axon, we loaded the nodes in each axonal skeleton into a unique KD-Tree that we queried with points spaced 1-μm apart on the dendritic tree. The number of these points that were within 5μm of a point in the axon’s KD-Tree proportionately determined the probability of shuffling a synapse between that axon and dendrite to that edge. For each dendritic tree and for each of its presynaptic axons, we recorded the list of dendrite edges proximal to the axon and the probability of choosing each edge. To proximally shuffle, we referenced this table to probabilistically choose which edge to position a synapse on a dendritic tree.

To shuffle synapses within a dendritic branch, we found the distal-to-proximal ordering of its *n* synapses and applied a random permutation sampled uniformly from the set of length-*n* permutations.

### Bayesian model of synaptic clustering

We represent the probability that a pair of axons clusters on a branch of dendrite with a Bernoulli-distributed random variable *p*_*c*_ parametrized with a Gamma-distributed random variable *β*. Concretely, for a dendritic tree *t*, a pair of axons *i* and *j*, and a set of branches *S*_*i,t*_ or *S*_*j,t*_ where axon *i* or *j* may synapse (table of proximities):

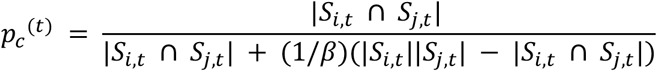

| *S*_*i,t*_ ∩*S*_*j,t*_ | counts the number of branches of tree *t* proximal to both axon *i* and *j* and | *S*_*i,t*_ || *S*_*j,t*_ | counts the total number of pairs of branches from *S*_*i,t*_ and *S*_*j,t*_ that axon *i* and *j* could choose to synapse onto. If β = 1, then 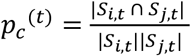, the probability that axons *i* and *j* cluster on a branch of tree *t* if they independently choose to synapse onto a branch in *S*_*i,t*_ and*S*_*j,t*_, respectively. As β → ∞,*p*_*c*_^(t)^ → 1 and as β → 0,*p*_*c*_ ^(t)^ → 0. Thus, β > 1 indicates attraction and β < 1 indicates repulsion. To extend this approach to larger groups of axons (*k*> 2), we modeled the first *k*− 1 axons as a single axon and the *K*-th axon, held-out from this group, as another axon. Given a dendritic tree *t*, we find the set of branches proximal to the *K*-th axon as usual (call this *S*_*K,t*_) and those proximal to the other *k*− 1 axons (call this *S*_*K*-1,*t*_). We repeat this process of finding proximal branches for all *k* choices of the held out axon, since we do not know *a priori* which axons belong to the same ensemble and which do not. Thus, we obtain *k* clustering probabilities for each neuron postsynaptic to the axon group.

To infer whether a pair is repelled from or attracted to the same dendritic branch given its observed clusters across a set of postsynaptic trees, we introduced two competing prior distributions for *β*, one with mean ½ and variance 1, truncated to the domain [0, 1], and the other with mean 2 and variance 1, truncated to the domain [1, ∞). Starting from a list of every dendritic tree postsynaptic to both axons *i* and *j*, we recorded a list of binary observations c*t* for whether the pair clustered on a branch of tree *t* and used the Python library PyMC (NUTS algorithm) [20] to optimize both models with this same list of observations. The likelihood that the attracting model and the repelling model reproduce these observations is obtained as a result of PyMC model optimization, and the ratio of the former to the latter yields a Bayes factor. There is a single Bayes factor for each pair of axons and *k* factors for a group of *k*> 2 axons, corresponding to the *k* choices of held-out axon for that group. We select the most extreme-valued of these *k* Bayes factors, which corresponds to the most likely clustering scenario. The mean of β’s posterior distribution for the more likely model given these observations yielded the mean clustering bias for that group of axons.

### Predicting synaptic clusters by cortical layer and axon group

We predicted the presence or absence of a synaptic cluster based on the inferred clustering biases. To predict whether an axon group clusters on one of its postsynaptic neurons, we calculated the axon-wise clustering probability *p*_*c*_ ^(*t*)^(*β*) based on the group’s inferred clustering bias β′, stochastically choosing whether the group clusters by sampling from a Bernoulli distribution with probability *p*_*c*_ ^(*t*)^(β′). The likelihood of the true observation (presence or absence of cluster) was calculated as the likelihood of a success or failure from the sampled Bernoulli distribution.

To estimate the clustering probability of a synaptic cluster from an axon pair based on cortical layer, we averaged the axon-wise clustering probabilities *p*_*c*_ ^(*t*)^(β) of every axon pair (*i,j*) from layers (*L*_*i*_,*L*_*j*_) that is proximal to a dendrite in layer *L*_*d*_. This average estimates the probability that an axon from *L*_*i*_ clusters with an axon from *L*_*j*_ on a dendrite in *L*_*d*_. We estimated the significance of this clustering probability by repeating the above process but assuming β = 1 for every pair and then calculating a paired-sample t-score for each triplet of cortical layers, measuring the extent to which the mean of the inferred clustering probabilities deviated from the mean of the baseline clustering probabilities.

To predict the presence or absence of a synaptic cluster based on layer-wise clustering probabilities, we extracted the cortical layers of the axon group and of the postsynaptic dendrite. If the group contains only two axons, then we directly use the layer-wise clustering probability as the probability of success in the Bernoulli distribution, sampling as before to stochastically choose whether the pair clusters on the dendrite. If the axon group contained more than two axons, then we calculate the probability that the held-out axon clustered with the rest of its group as the product of *k* − 1 layer-wise clustering probabilities that describe the probability that an axon from the held-out axon’s layer clusters with other axons from layers of the remaining group onto a dendrite from the same layer as the postsynaptic dendrite. This product is used as the probability of success in the Bernoulli distribution that we sample to predict whether the group clusters on the dendrite.

### Bayesian model of order preference

For each axonal group that clusters its synapses onto branches of at least two different dendritic trees, we fit a Bayesian model that explains each ordering of its synaptic cluster as samples from a Mallows distribution. Its prior dispersion is drawn from a uniform distribution between 0 and 1 and its prior central permutation is drawn from a Mallows distribution whose dispersion is 1 and central permutation is the identity (without loss of generality). As done for the Bayesian models of synaptic clustering, we used PyMC’s NUTS algorithm to optimize this ordering model and extract the resulting likelihood of the observed orderings in each cluster of a particular axonal group.

To determine whether an axonal group prefers to synapse in a particular order, we compared its likelihood under the dispersion-optimized model to its likelihood under an alternative model where the dispersion’s prior distribution is fixed at 1. With its fixed dispersion, this maximum-dispersion model’s central permutation can be arbitrarily chosen as the identity permutation and the likelihood of the data given by a uniform distribution with probability 1/*k*! for all possible orderings. We calculated the Bayes factor for the dispersion-optimized model to the maximum-dispersion model in the same manner as for the Bayesian models of attraction to repulsion.

### Distal preference

To estimate a *distal preference* for each axon in an axodendritic permutation, we use the Bradley-Terry model of pairwise comparisons [14]. To find the probability *P*(*i distal to j*) = 1 − *P*(*i proximal to j*) of axon *i* synapsing distal to axon *j*, we use their axodendritic permutation’s inferred dispersion to calculate the probability of every permutation and marginalize across those for which axon *i* is indeed distal to axon *j*. Given *P*(*i distal to j*) for all *j*, the Bradley-Terry model infers a *latent score* for axon *i* via maximum-likelihood estimation. Normalizing these scores to [0, 1] yields axon *i*’s most likely distal preference. Given these inferred distal preferences, we use the Plackett-Luce model of ranking *N* > 2 items to determine a distribution over all distal-to-proximal permutations of this group of axons [15,16].

We estimate the distal preference of axons from a given layer synapsing onto dendrites in a given layer as follows. First, we sum the subset of probabilities of permutations that preserve the order of pairs of axons from different cortical layers across the Mallows distribution fitted to an axodendritic permutation to obtain an order probability. Next, we find the mean of these probabilities across all axodendritic permutations clustered on dendrites in a given layer to estimate the probability that an axon from the first layer synapses distal to an axon from the second layer on a dendrite in the third layer. Finally, we use the Bradley-Terry model to infer a distal-preference of this pair of layers and the Plackett-Luce model to determine a distribution over all distal-to-proximal permutations of the cortical layers.

### Predicting synaptic orderings by axon group, axon pairings, and layer pairings

To predict synaptic orderings by axon group, we used the inferred dispersion and central permutation of each axodendritic permutation to generate a Mallows distribution over their possible orderings. Alternatively, we use axon pairings or layer pairings to compute a distribution over the group’s synaptic orderings as before: inferring a latent distal preference via maximum-likelihood estimation in the Bradley-Terry model and computing a distribution over permutations via the Plackett-Luce model. The predicted ordering for a group’s cluster was the most probable ordering under the computed distribution, with ties broken randomly. The likelihood of a group’s true synaptic ordering was taken as the probability of that ordering under the computed distribution.

**Acknowledgments**

We thank F. Collman for technical assistance with the MICrONS Cubic Millimeter dataset and A. Tolias for his advice and support. We also thank members of the Tolias Lab, Z. Ding and S. Papadopulous, and of the Boahen Lab, G. Weintraut and N. Reidman, for their helpful technical discussions throughout the study.

## Supplementary Figures

**Supplementary Figure 1:**
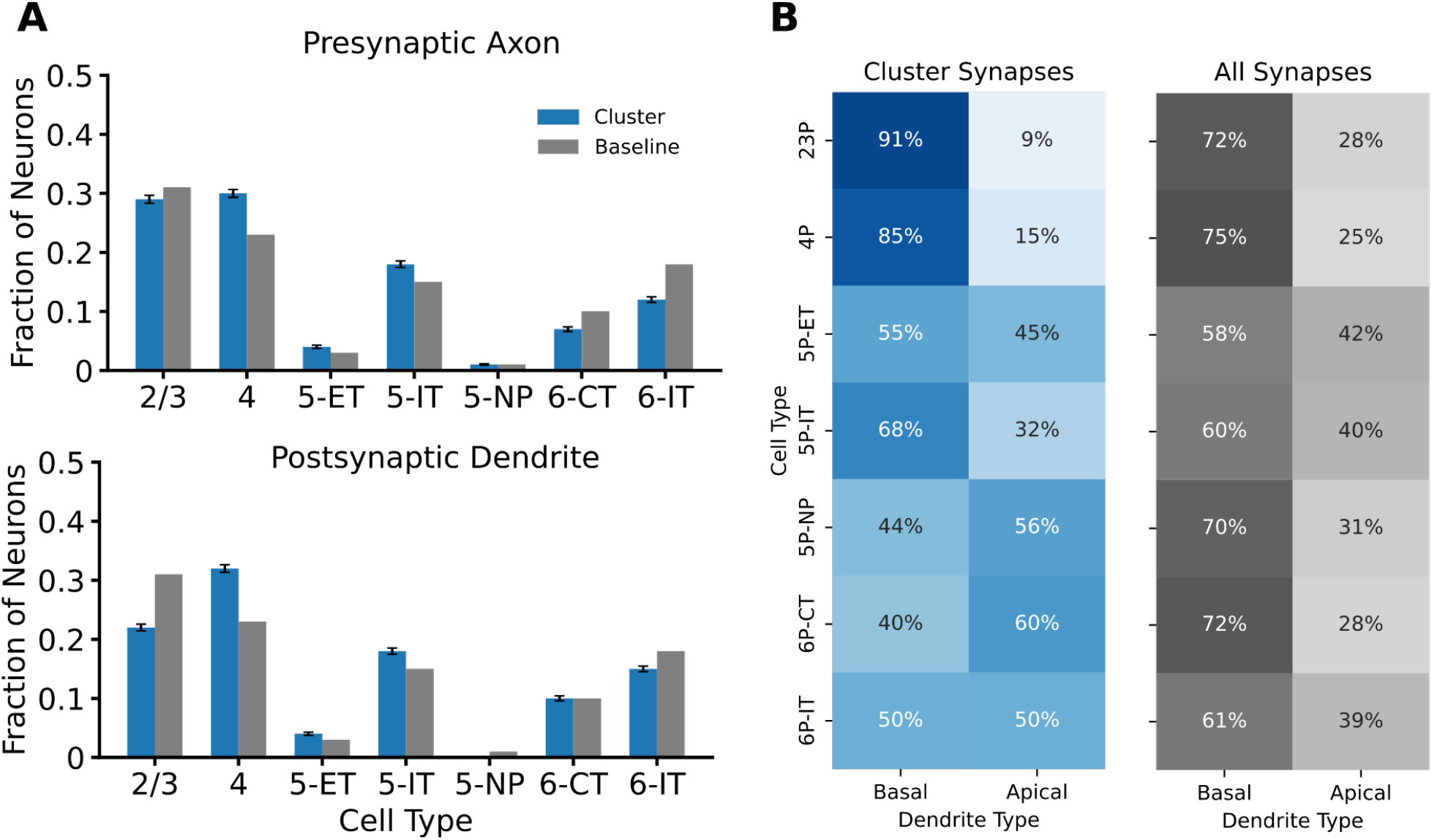
Synaptic clusters found on basal and apical dendrites of various pyramidal-cell types. A – Fraction of axons and dendrites: Axons that form (top) and dendrites that receive (bottom) synaptic clusters compared to the fraction of cells of that type (gray). Pyramidal cells in Layers 5 and 6 are subdivided into extratelencephalic (ET), intratelencephalic (IT), corticothalamic (CT), and near-projecting (NP). Bar, 95% credible interval. B – Distribution of synapses within clusters (left) by cell type and dendritic compartment compared to the distribution of all synapses (right).

**Supplementary Figure 2:**
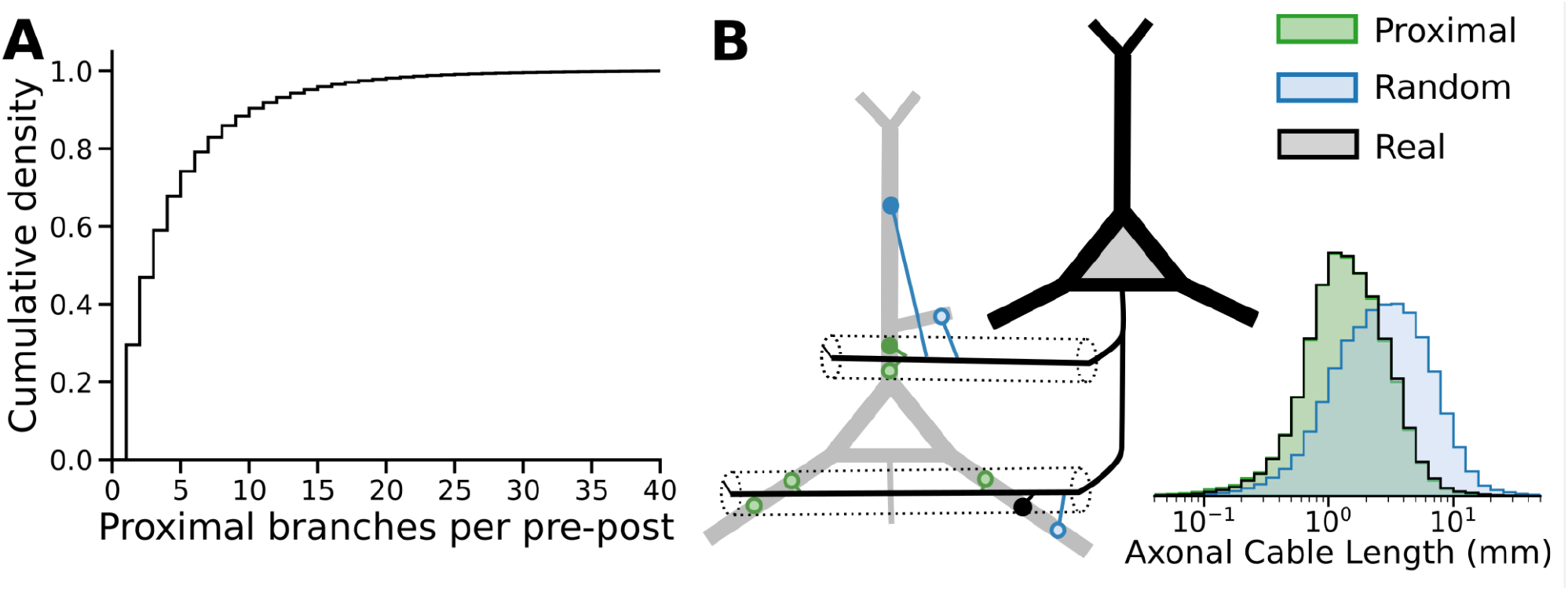
Dendritic branches proximal to a presynaptic axon. A – Cumulative distribution: Across all pairs of pyramidal cells connected with at least one synapse, the median number of dendritic branches proximal to a presynaptic axon is 3. B – Random and proximal shuffles: Each of 4 million synapses connecting 48 thousand pyramidal cells is shuffled to a random (*blue*) or proximal (*green*) site on its postsynaptic dendritic tree. The proximal site is within 5 μm of the presynaptic axonal arbor and thus preserves the connectome’s axon-length distribution (*black outline*, 2-sample Kolmogorov-Smirnov (K-S) statistic = 0.0055; double-sided *P* = 0.37), whereas the random site triples the median axon-length (K-S = 0.31; *P* < 10^−16^).

**Supplementary Figure 3:**
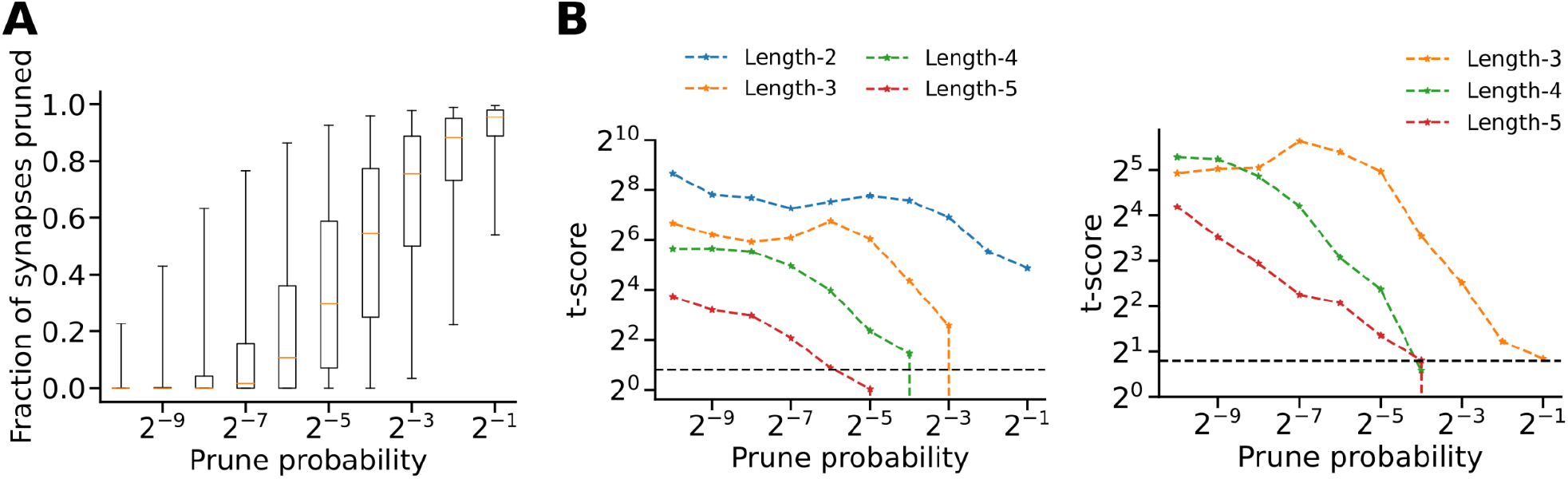
Synaptic cluster repeats in probabilistically unmerged axons. A – Probabilistically unmerging and discarding distal axonal branches prunes synapses from the connectome (*n* = 50). B – Significance of unordered and ordered synaptic clusters in real vs. proximally-shuffled pruned connectomes (*n* = 50): *Left*, Unordered clusters of 5, 4, and 3 axons drop below significance (*P <* 0.05, *black line*; degrees-of-freedom = 49) at a prune probability above 1/64, 1/16, and 1/8, corresponding to a connectome where 21%, 51%, and 67% of synapses are pruned on average. *Right*, Ordered clusters of 5, 4, and 3 axons drop below significance at a prune probability above 1/32, and 1/2, corresponding to a connectome where 35% and 91% of synapses are pruned on average.

## Notes

### Competing Interest Statement

The authors have declared no competing interest.

